# Investigating experimental storage methodologies for the understudied intermediate recalcitrant seed of Northern Wild Rice (*Zizania palustris* L.)

**DOI:** 10.1101/2023.08.03.551837

**Authors:** A. Mickelson, L. McGilp, J. Kimball

## Abstract

The maintenance of plant genetic diversity is an essential target for conservation and breeding efforts. However, the ability to store seed varies between species with those that are more easily stored being overrepresented in seed banks. Northern Wild Rice (NWR; *Zizania palustris*) has intermediately recalcitrant seed which, along with its dormancy period, makes it more challenging to store long term. This study evaluated alternate storage treatments, including water changes, aeration, and the shift from submerged to moist seed, to extend the longevity of NWR seed in storage compared to current best management practices. Monthly water changes were the most effective storage treatment, maintaining greater than ∼ 50% viability for over 28 months. There was a negative correlation found between aerated treatments with high dissolved oxygen and seed viability. Submerged to moist storage was only effective for ∼16 months. Control treatments maintained relatively high viability (≥ 58%) through 21 months of storage. However, by 28 months, monthly water change seed had significantly higher viability (57.6%) compared to either the standard control (37.2%) or the bucket control (28.9%), suggesting that this treatment is more efficacious than standard storage conditions for maintaining seed viability of NWR.

## 1 Introduction

As plant diversity is eroding on a global scale at alarming rates, concerted efforts to maintain and conserve that diversity have focused on the ex-situ storage of germplasm outside of species’ native habitats (Parmesan and Hanley 2015; Wyse *et al*. 2018; Breman *et al*. 2021; Pence *et al*. 2022). Today, over 100,000 species are conserved via ex-situ methods (Breman *et al*. 2021), including a variety of sociologically and economically important species as well as those threatened by habitat alterations, new diseases and pests, and extinction (Liu *et al*. 2018; Wyse *et al*. 2018; Walters and Pence 2021; Pence *et al*. 2022; Fernández *et al*. 2023). This germplasm can be preserved, while also allowing for its use in agriculture, restoration projects, species reintroduction, biological control, breeding, and research (Liu *et al*. 2018; Walters and Pence 2021). The majority of long-term ex-situ germplasm storage occurs in roughly 1,750 seed or gene banks, which provide freezer or cryogenic storage of seed and other tissues for around 50-60 thousand taxa (Walters and Pence 2021).

Species vary in their ability to be stored, particularly as seed, and estimates suggest that close to half of all plant species may not be storable by these conventional methods (Li and Pritchard 2009; Wyse *et al*. 2018; Hay and Sershen 2021; Walters and Pence 2021; Fernández *et al*. 2023). Orthodox seed, which is desiccation tolerant, for example, can be dried and frozen, halting metabolic functions. These seeds are often storable for decades to centuries and represent the majority of seed currently present in seed banks (Walters *et al*. 2005, 2013; Walters 2015; Liu *et al*. 2018). Recalcitrant seed or desiccation sensitive seed, however, are more challenging to store long-term and are not amenable to traditional storage techniques employed by seed banks (Wyse *et al*. 2018; Walters and Pence 2021). Instead, for many recalcitrant species, the only long-term storage method is through the cryopreservation of their embryos (Walters *et al*. 2001; Berjak and Pammenter 2008). However, embryo extraction and cryogenic freezing are not currently feasible on a large scale for the majority of recalcitrant species. Aside from cryopreservation, the general best practice for the storage of recalcitrant seed is to store seeds as close to their shedding water content as possible and at as low a temperature as can be tolerated (Berjak and Pammenter 2008; Pammenter *et al*. 2014). This is termed hydrated storage and is considered effective only in the short to medium term, particularly for chilling sensitive species, due to microbial proliferation and the eventual germination of seed (Berjak and Pammenter 2001, 2008, 2013; Pammenter *et al*. 2014). Hydrated storage may extend seed viability anywhere from days to months for chilling sensitive species, or up to 1-2 years for some temperate species (Pammenter *et al*. 2014).

Seed recalcitrance is found to some degree in ∼8% of all seeded plant species, ∼1.4% of species in the Poaceae family, and has developed multiple times across phylogenetically distinct taxa (Dickie and Pritchard 2002; Wyse and Dickie 2017). The majority of recalcitrant seeded plants are non-pioneer trees from tropical or aquatic environments, where high levels of plant diversity as well as threatened species can be found, making the preservation of recalcitrant species from these regions an important goal for the conservation of biodiversity (Tweddle *et al*. 2003; Berjak and Pammenter 2008; Wyse and Dickie 2017; Wyse *et al*. 2018). Among recalcitrant species, there is a wide range of variability in the extent of desiccation tolerance, critical moisture content, and storage longevity, in some cases even within a single genus (Berjak and Pammenter 2008, 2013; Pammenter *et al*. 2014; Umarani and Groot 2023).

Like many recalcitrant species, the ex-situ storage of Northern Wild Rice (NWR; *Zizania palustris* L) is limited using conventional storage methods and is currently not stored in seed banks due to its unique seed physiology. NWR is an open-pollinated, annual, aquatic plant in the *Poaceae* family. NWR grows naturally in slow moving lakes and streams, primarily in the Great Lakes region of North America, and is cultivated in Minnesota and California as a small commodity crop in man-made dike-lined paddies (Oelke and Porter 2016; McGilp *et al*. 2023). The seed of NWR is intermediately recalcitrant, meaning that it falls somewhere between orthodox and recalcitrance in terms of its response to desiccation, storage temperature, and critical moisture contents (Ellis *et al*. 1990; Ellis 1991; Walters 2015), and results in unique challenges for storage longevity (Probert and Longley 1989; Ellis 1991; Kovach and Bradford 1992a; b; Walters 2015; McGilp *et al*. 2020). The tolerance to desiccation in NWR can be affected by a number of factors including temperature, drying rate, and degree of metabolic activity at the time of storage (Probert and Longley 1989; Kovach and Bradford 1992a; Berjak and Pammenter 2008). To accommodate the desiccation sensitivity of NWR, the current best practice for storing seed includes submerging seed, after post-harvest processing, in sealable plastic bags and moving them to a dark, 1-3°C cooler (Kovach and Bradford 1992b; McGilp *et al*. 2020). However, under these storage conditions seed only remains viable for 1-2 years (Grombacher *et al*. 1997). This means that the breeding and research of NWR requires that all germplasm lines be grown yearly, taking up valuable resources.

Northern Wild Rice also has a seed dormancy period of ∼3-6 months before reaching maximum germination, though seed may reach around 50% germination by ∼8 weeks in storage (Cardwell *et al*. 1978; McGilp *et al*. 2022). To break dormancy in cultivated NWR, seed must be stratified under cool (1-3 °C), submerged, and dark conditions (Simpson 1966; Kovach and Bradford 1992b; Grombacher *et al*. 1997; McGilp *et al*. 2020). As dormancy begins to break during storage, seed viability tends to rapidly decrease and is accompanied by the growth of both bacteria and fungi (Berjak and Pammenter 2008). It has been hypothesized that dormancy in NWR seed is either physiological or morphophysiological, requiring cold stratification and potentially, an interaction with gibberellic acid to germinate (Baskin and Baskin 2004; McGilp *et al*. 2022). There are many knowledge gaps regarding the interplay between seed recalcitrance and dormancy in NWR and research is ongoing. However, both conditions complicate the storage of NWR seed.

Several studies have looked at alternate storage methods for NWR seed including variations in temperature, seed moisture content, and environmental moisture conditions, but they have shown limited long-term utility (Oelke and Stanwood 1988; Kovach and Bradford 1992a; McGilp *et al*. 2020). Anecdotally, seed stored by MN cultivated NWR growers in large bags placed into drainage ditches, appears to maintain viability, and have improved germinability relative to breeding program seed. While it is unclear what factor contributes most to this difference, it is hypothesized that water movement and the resulting increase in oxygen within the water could be involved. The effects of dissolved oxygen (DO) on various stages of plant growth have been documented over time. The presence of oxygen has been shown to increase germination in some aquatic plants such as *Vallisneria americana* (Jarvis and Moore 2008), while in others, such as *Zizania Texan*, too high of DO can decrease germination (Power and Fonteyn 1995). Additionally, a higher DO appears to improve the germination and growth of wheat (Zhu *et al*. 2021). However, no such studies have been conducted to determine the effect of DO on the long-term storage of recalcitrant seed. This study aimed to fill in this knowledge gap and to test the efficacy of alternate storage treatments for extending the viability of NWR seed in storage. The conditions tested in this study included the effects of DO and water changes on the longevity of seed. In addition, due to previous observations that seed viability may be maintained following dormancy break without seed submersion in water (Supp figure 1), the shift from submerged to non-submerged storage was also tested in this study.

## 2 Materials and methods

### Plant material

The current MN industry standard, cv. “Itasca-C20”, was selected for these experiments. Plants were grown in open-pollinated 12 x 24 m dike-lined paddies at the North Central Research and Outreach Center (NCROC) in Grand Rapids, MN. Seed was harvested from ∼20 individual plants in bulk in September of 2020 and placed immediately into treatment bags, as described below.

### Experimental Setup

To assess alternative ex-situ storage methods for NWR seed, five unique seed storage methodologies, along with two controls, were evaluated in this study. These treatments included non-aerated and aerated methods, weekly and monthly water changes, a combination of the two, and a submerged to moist treatment based on previous anecdotal observations of its efficacy. The control treatments consisted of a standard control (SC) with seed stored in the dark in sealed plastic 30.5 x 30.5 cm bags with 0.7 liters of reverse osmosis (RO) water, which is consistent with current best practices for storing NWR seed (Simpson 1966; Cardwell *et al*. 1978), and a bucket control (BC) with seed stored in 20.3 x 30.5 cm mesh bags and placed into a 24.6 liter brewers bucket with 4.3 liters of RO water. The BC was used to determine whether any differences would exist between a mesh bag with water and the SC in plastic bags. Weekly and monthly water change (WWC and MWC, respectively) treatments were stored in the same fashion as the bucket control but with water drained through the spigot and replaced with cooled RO water of the same volume on a weekly or monthly basis, respectively. To test the impact of DO levels on NWR seed viability in ex-situ storage, a circular pond air stone (Aquascape 75001, St. Charles, IL) was used for the aeration (ABC) and aeration with weekly water change (AWC) treatments, which were stored identically to the BC. For the two-stage storage (2SS) treatment seed was stored like the BC until 21 weeks of storage, then drained and left damp for the remainder of the experiment.

A total of 21 brewer’s buckets, each representing one of three replications per treatment, were organized in a dark 1-3°C cooler as a completely randomized design. Each bucket contained 6 plastic or mesh bags, depending on the treatment, filled with ∼1950 Itasca-C20 seeds to test germination over six individual time points. To eliminate air flow in the brewer’s buckets, the holes in all buckets, except those used for aeration treatments, were plugged. The temperature of the cooler was monitored throughout the study using a HOBO logger (Onset MX2302A, Bourne, MA). A DO meter (Ecosense ODO200M, Berlin, Germany) was also placed in one replication of each of the treatments to monitor DO changes across the storage period.

In total, the experimental treatments were evaluated 5 times over the course of a 28-month period, starting immediately after harvest, apart from two treatments requiring weekly water changes, WWC and AWC. These two treatments were only measured through 21 months in storage due to the intense labor required and lack of resources. The final (6th) time point of the experiment was dropped in this study due to a cooler malfunction, which froze the entire experiment.

### Dissolved oxygen monitoring

Dissolved oxygen was measured weekly for the AWC and WWC treatments and monthly for all other treatments. For the 2SS treatment, DO was only measured through 21 weeks, as the water was removed following this time point. During the time of DO measurements, the room was kept dark to decrease the exposure of seeds to light. To measure DO, the buckets were individually removed from the cooler and the probe was placed below the bags within the bucket and moved around briefly. The DO measurement was taken after about 2 minutes to ensure that the probe had reached equilibrium.

### Germination and Tetrazolium testing

Germination data was collected at 4, 9, 16, 21, and 28 months in storage, beginning in January of 2020 and ending in January of 2023. For each timepoint one bag of seed was removed at random from each brewer’s bucket. When seed bags were removed at each time point, water was also removed to maintain a 3:1 ratio of water to seed. From each bag, 20 seeds were moved to a new 10 x 10 cm plastic bag with 20 ml of RO water and placed in a growth chamber set to 15-hour days and 27°C. After 7 days the water in each bag was emptied and replaced. After 14 days, the number of germinated seeds, defined as coleoptile growth past the end of the seed, was recorded for each rep of each treatment. Non-germinated seeds were then stained with 2,3,5-Triphenyltetrazolium chloride (TZ; Carolina Biological Supply, Burlington, NC) by cutting the seeds lengthwise to expose the embryo, placing them into plastic segmented storage containers with enough 0.2% TZ to fully submerge seed, and after two hours, recording the number of seeds with embryos that had stained red. Percent viable seed was calculated as 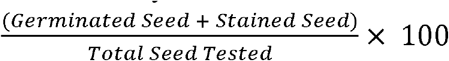.

### Data Analysis

Analysis of the data was conducted in Rstudio, version 2023.03.0 (Rstudio-team 2023). Data were run through standard diagnostic plots, then analyses of variance were conducted with percent viable seed and dissolved oxygen level as response variables. Significant factors were then compared using Tukey’s HSD, calculated using the agricolae package, version 1.3-5 (Mendiburu 2021). Contrasts were calculated using the Hmisc package, version 5.0-1 (Harrel 2023). All plots were made using the ggplot2 package, version 3.4.1 (Wikham 2016).

## 3 Results

### Dissolved oxygen

To assess the effect of aeration on the viability of NWR seed in storage, DO measurements were taken throughout the course of this experiment. The DO was measured weekly or monthly for 29 months, dependent upon treatment, except for the 2SS treatment, which was drained of water at 22 weeks, to assess the effect of reducing water post-dormancy break. The AWC and WWC treatments ended at 22 months due to their laborious nature along with their low germination rates after 22 months. An ANOVA of this data found significant differences between treatments, months in storage, and the interaction of the two (Table 1). The two control treatments, BC and SC, differing only in the material of the storage bags, maintained a relatively constant DO, never going outside of the 0-2 mg/L range (Figure 1). There were no significant differences between these treatments. The two aerated treatments, ABC and AWC, maintained significantly higher DO levels throughout the course of the study (10.8-13.3 mg/L and 11.5-13.3 mg/L, respectively) than any of the other treatments or the controls. Both aerated treatments had wider ranges of DO than either control. Compared to the controls, the DO of the WWC and MWC treatments, which had water changes weekly or monthly, was slightly higher on average. The MWC remained within 1 mg/L of the controls for the full duration of the study. The WWC was more variable but overall maintained higher DO than both of the controls and the MWC treatment. There were DO peaks (∼2.8 mg/L) at around 6 months and 22 months for the WWC, with declining DO between them. The AWC treatment had oxygen introduced both through weekly water changes and by aeration of the water. After around 3 months of storage, this treatment retained the highest DO through 22 months, when measurements ceased. Additionally, the AWC treatment appears to vary in a similar pattern to both the ABC and the WWC treatments, with peaks around 6 and 22 months. However, the ABC treatment, while only fluctuating over a small range, appears to trend downward over time. It is unclear if the AWC and WWC, the other two most heavily fluctuating treatments, would also have trended downwards, as the measurements were ended prior to this larger drop in the ABC treatment. The 2SS treatment was only submerged, and therefore had DO measured, through 5 months. The DO of this treatment directly mirrored that of the BC during this time, which is expected, as they were stored identically until 5 months of storage. Overall, more variability was seen between the monthly DO ratings of treatments that included more perturbation of the seed, including both aerated treatments and the WWC treatment (Figure 1).

**Table 1.** Analysis of Variance (ANOVA) of dissolved oxygen (DO) levels (mg/L) collected over the span of 28 months across seven storage treatments of Northern Wild Rice (NWR; *Zizania palustris*).

|  | df | Sum Squares | Mean Squares | F-value | Pr(>F) |
| --- | --- | --- | --- | --- | --- |
| **Timepoints** | 28 | 150.38 | 5.37 | 40.02 | < 0.001 |
| **Treatments** | 6 | 13022.22 | 2170.37 | 16173.20 | < 0.001 |
| **Timepoints:Treatments** | 130 | 48.62 | 0.37 | 2.79 | < 0.001 |
| **Residuals** | 330 | 44.28 | 0.13 |  |  |

**Figure 1.**
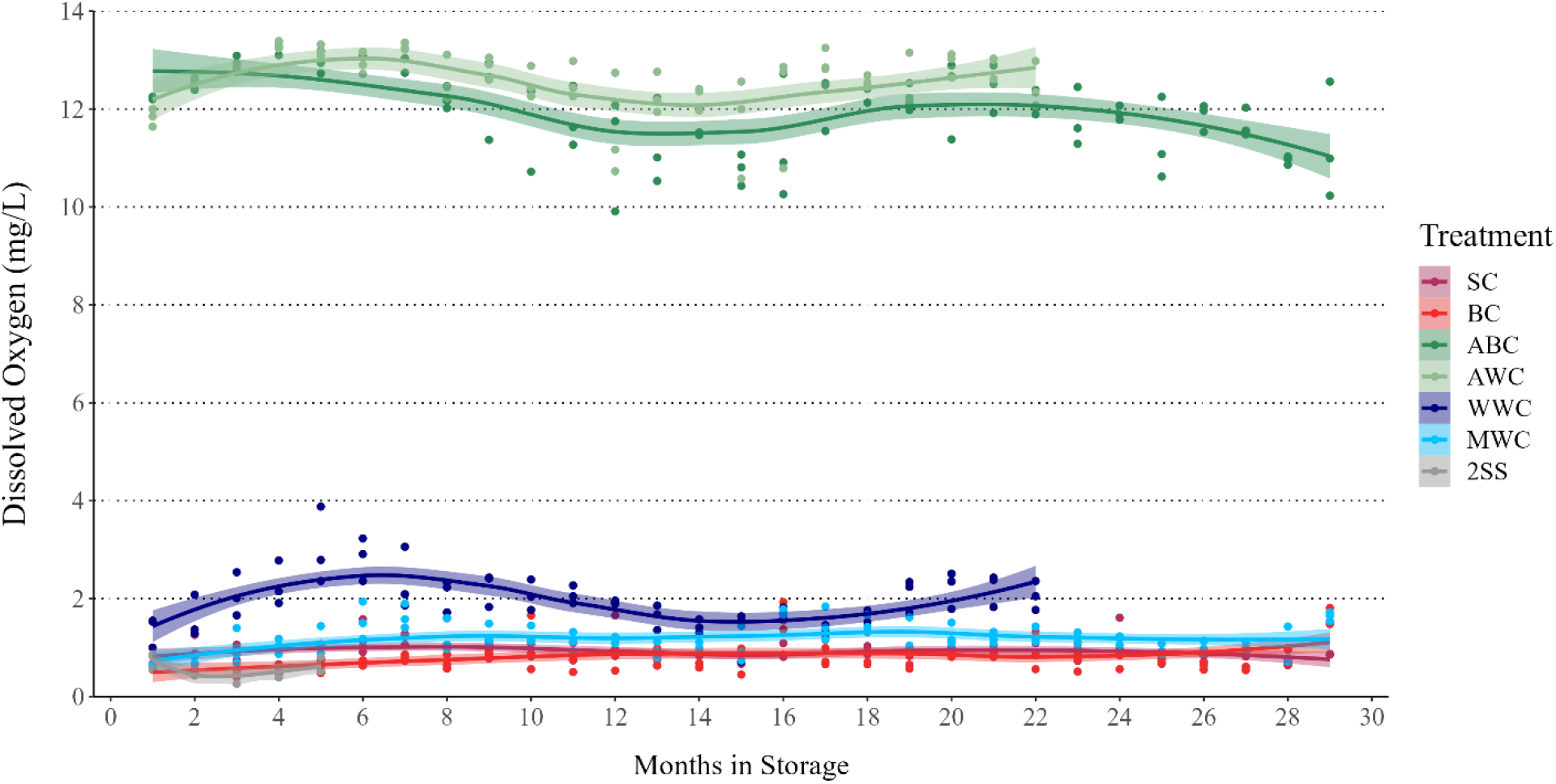
Dissolved oxygen levels (mg/L) in standard and experimental Northern Wild Rice (NWR; *Zizania palustris* L.) storage conditions monitored daily across a 28-month time period.

### Seed Viability

Seed viability was tracked in this experiment to determine the most efficacious storage treatments for extending seed longevity in NWR. Germination and tetrazolium staining data were taken for each treatment at 4, 9, 16, 21, and 28 months and used to calculate percent seed viability. The AWC and WWC treatments were only measured to 21 months of storage due to low germination and the laborious nature of these treatments. An ANOVA found significant differences in seed viability between treatments, months in storage, and their interaction (Table 2). There were no significant differences in viability between the two control treatments, SC and BC, over the course of the study. Both controls matched or exceeded the viability of all other treatments until 28 months in storage, at which time the viability dropped steeply to 37.2% and 28.9%, respectively. The largest initial drop in viability occurred at 9 months of storage in the ABC (−31%) and AWC (−17%) treatments, which were both aerated treatments with high DO. Following this initial drop, the aerated treatments had the lowest seed viability ratings for all timepoints between 9 and 22 months of storage, falling below 50% by 16 months of storage (Figure 2). The water change treatments, WWC and MWC, maintained similar viability to the controls until around 16 months in storage, at which point there was a large drop in viability for the WWC treatment (−33.6% from 9 to 16 months), as well as an increase in variability between reps of the treatment. Unlike the WWC treatment, the MWC, which was disturbed less often, maintained high viability through the remainder of the experiment, exceeding that of both controls by 28 months in storage (MWC 57.6 %; SC 37.2 %; BC 28.9 %). The 2SS treatment was based upon prior observations that viability could be maintained when seed was only submerged through the duration of dormancy break (Supplemental figure 1). The 2SS treatment was submerged through 5 months of storage and maintained as good or better viability than both controls through 16 months in storage. However, by 21 months the seed viability had dropped precipitously from 75.9% at 16 months to 26.4% at 21 months. By 28 months the 2SS had the lowest viability (11.9%) of any treatment or control. At 21 months, the last time point that contained all the treatments, the SC, BC, and MWC had the highest viability, with no significant differences between them (66.2 %, 73.0%, 62.3%, respectively). When averaged across treatment months, seed viability was highest in the MWC (77.2%), BC (76.0%), SC (75.3%), and WWC (67.0%) treatments. When averaged across treatments, seed viability consistently dropped between each timepoint, starting at 96.7% at 4 months in storage and ending at 35.8% at 28 months. Additionally, the longer the seed was in storage the greater the chance for variation between reps of each treatment.

**Table 2.** Analysis of Variance (ANOVA) of Northern Wild Rice (NWR; *Zizania palustris*) seed viability (%) collected over the span of 28 months (5 timepoints) across seven storage treatments.

|  | df | Sum Squares | Mean Squares | F-value | Pr(>F) |
| --- | --- | --- | --- | --- | --- |
| **Timepoints** | 4 | 59482.55 | 14870.64 | 78.35 | < 0.001 |
| **Treatments** | 6 | 11046.12 | 1841.02 | 9.70 | < 0.001 |
| **Timepoints:Treatments** | 22 | 9975.06 | 453.41 | 2.39 | < 0.001 |
| **Residuals** | 66 | 12527.23 | 189.81 |  |  |

**Figure 2.**
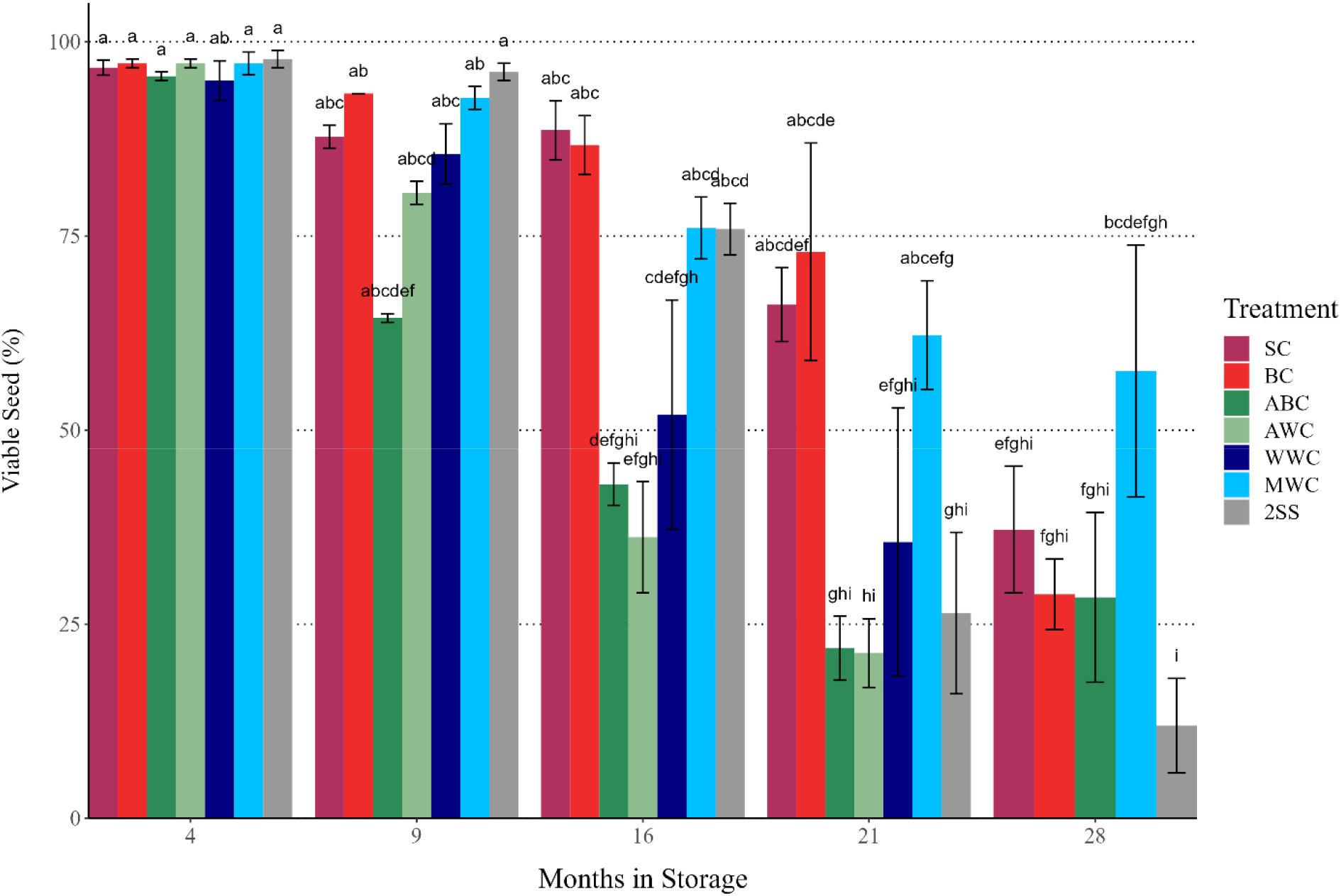
Adjusted means for % viable seed of Northern Wild Rice (NWR; *Z. palustris*) over the course of 28 months in seven different storage treatments including a standard control (SC); bucket control (BC); aerated bucket control (ABC); aerated weekly water change (AWC); weekly water change (WWC); monthly water change (MWC); and a two-phase seed storage (2SS) methodology. Tukey’s significance values were calculated across all treatments and time points.

## 4 Discussion

### Northern Wild Rice’s intermediate seed recalcitrance is understudied and a serious challenge for preservation and breeding efforts

Improving the ex-situ seed banking of plant species that are not amenable to current storage methods is crucially important for preserving present day biodiversity. Ex-situ conservation is very limited for both wild populations and cultivated NWR, as there is no current best management practice that can maintain NWR seed viability for longer than ∼1-1.5 years. Like many plant species worldwide, the genetic diversity of NWR is eroding rapidly in the species centers of origin and diversity in the Great Lakes region of North America (Norrgard 2006; Hansen 2008) and the species appears to be slowly migrating northward (Terrell *et al*. 1997). This erosion is due in part to shifts in water levels, pollution from shoreline activities, mining, and the effects of climate change (Hansen 2008). The species has also recently been red listed by the International Union for Conservation of Nature (Maiz-Tome 2016). The recalcitrant nature of NWR seed, in combination with its dormancy, also presents a challenge for cultivated NWR breeding programs, requiring yearly maintenance of all germplasm, which is labor and resource intensive. The laborious nature of germplasm maintenance for NWR under current best management practices puts many of the services this species provides at risk.

Most of the work in seed recalcitrance has been done with recalcitrant tropical tree species, where hydrated storage or the storage of seeds at high relative humidity, has been used to moderately extend the longevity of some recalcitrant species (Pammenter *et al*. 1994, 1995; Berjak and Pammenter 2008; De Vitis *et al*. 2020). Studies evaluating intermediate recalcitrance are rare in the literature as is submerging recalcitrant seed during storage, only seemingly being used for NWR (Kovach and Bradford 1992a; McGilp *et al*. 2020) and a handful of seagrass (*Zostera*) species (Conacher *et al*. 1994; Yue *et al*. 2019; Infante-Izquierdo *et al*. 2023). Current research has determined that NWR seed should be submerged and stored at cool temperatures to extend seed longevity and allow for dormancy break (Simpson 1966; Cardwell *et al*. 1978; Kovach and Bradford 1992b; McGilp *et al*. 2020).

### High dissolved oxygen levels are tied to decreased Northern Wild Rice seed viability in submerged storage

There has been speculation that oxygen levels may be important for the viability of cultivated NWR seed, as seed from growers, which is stored in areas with flowing water, anecdotally maintains viability for longer periods. Although there is limited research in this area, aerobic submerged storage conditions have been shown to extend the seed longevity of seagrass, *Zostera capricorni* (Conacher *et al*. 1994). Results from this study, however, demonstrated that aeration and by extension high DO appears to have a negative effect on the storage longevity of NWR. As quickly as 9 months in storage, the aeration (ABC) and aeration with weekly water change (AWC) treatments, which had significantly higher DO levels, were the first of the storage methods to decline in viability. High oxygen levels have been shown to shorten the viability of orthodox seed from a number of species, resulting in recommendations to control oxygen levels in storage (Roberts and Abdalla 1968; Borisjuk and Rolletschek 2009; Groot *et al*. 2012).

While it is currently unclear how high DO levels led to declines in NWR seed viability, it is possible that high DO could be linked to an increase in respiration of both seeds and microbes. High levels of oxygen have been shown to increase the rate of metabolic processes in seeds, ultimately increasing ATP production and cellular respiration (Dongen *et al*. 2004; Borisjuk and Rolletschek 2009; Gupta *et al*. 2009). It has also been demonstrated that in recalcitrant seeds of *Quercus robur* and *Castanea sativa*, respiration rates increase in the presence of excess oxygen (Greggains *et al*. 2000). In recalcitrant seed storage, low temperatures are often used to delay germination, in part by decreasing the metabolic rate of the seeds (Berjak and Pammenter 2008; Szuba *et al*. 2022). It is possible that excess DO overcame this deceleration of metabolic activity, perhaps by creating a stress environment that triggered early dormancy break and subsequent germination, or more generally by increasing the respiration rate of the seeds. Higher oxygen levels also appear to increase microbial activity (Visnovsky *et al*. 2008; Umarani and Groot 2023). While microbial activity was not measured in this study, microbial growth has been cited as a major challenge for the storage of NWR, as well as other recalcitrant species such as *Q. robur* and *Avicennia marina* (Mycock and Berjak 1990; Finch-Savage *et al*. 2003; Berjak and Pammenter 2013).

It is also possible that the high availability of oxygen led to the increased production of reactive oxygen species (ROS). The production of ROS has been demonstrated to increase over time during the moist storage of recalcitrant seeds from *Q. robur* and *C. sativa*, resulting in abnormal seedlings, though this appeared to be independent of temperature and oxygen level (Greggains *et al*. 2000). Additionally, seeds of the recalcitrant species *Trichilia dregeana* displayed an increase in ROS production during hydrated storage, peaking at 4 months, with rapid viability loss occurring starting at 6 months (Moothoo-Padayachie *et al*. 2018). However, no information is available to link this increase of ROS production to changes in oxygen levels during the storage of recalcitrant seed. In addition, no studies of the production of ROS during storage of recalcitrant seed have been conducted under conditions of submerged storage.

### Northern Wild Rice seed viability in submerged storage is improved with monthly water changes

This study demonstrated that a new storage methodology, using monthly water changes, is effective in maintaining NWR seed viability above 50 % for more than 2 years. Comparatively, both controls had less than 37 % seed viability on average by the final time point. Although monthly water changes benefited seed viability across the full study timeline, weekly water changes caused a marked decrease in viability by 16 months of storage (85.6% at 9 months, 52.0% at 16 months). The WWC treatment fared only slightly better than the two aerated treatments by 21 months, though this difference was not significant. This suggests that the frequency of water changes during storage can greatly impact the success of this strategy for the maintenance of seed viability. It is possible that oxygen levels may account for the difference between the MWC and WWC treatments, since the DO of the WWC treatment was slightly higher than that of the SC, BC, and MWC. However, the DO was still well below that of the aerated treatments. Alternatively, as the WWC seeds were taken out of storage for water changes and measurements on a more frequent basis, it is possible that its decreased viability relative to the MWC was due to increased exposure to germination cues such as light or temperature fluctuation, despite our best efforts. It is also possible that the success of the MWC is related to the decrease in microbial growth during storage. However, since microbial growth was not measured during this study, more research is needed to validate the hypothesis that its growth was less in water change treatments, and to determine the effect of the frequency of water changes on its growth.

### Future research needs for improving ex situ storage methods for Northern Wild Rice

The intermediate recalcitrance of NWR is understudied and our knowledge of the physiological, genetic, and molecular changes that occur to NWR seed during submerged storage is limited. In addition, the relationship between the dormancy and intermediate recalcitrance of NWR is not well understood. The loss of viability of recalcitrant seed in storage is thought to be associated with one or a combination of mechanical damage, metabolically induced damage, or denaturing of macromolecules (Umarani *et al*. 2015). Unlike orthodox seed, recalcitrant seed remains metabolically active following seed shed (Berjak and Pammenter 2008; Pammenter *et al*. 2014), which often coincides with shorter seed viability in storage (Kibinza *et al*. 2006; Bai *et al*. 2011). This study demonstrated that water changes increase the storage longevity of NWR seed. However, water changes for all stored seed is a laborious undertaking. It may be possible to reduce the labor required by beginning to change water after 9 months in storage, as no differences were evident between water change treatments and the controls up to this point.

However, a follow-up study would be required to validate this hypothesis. Future studies of seed storage methodology should consider the goal of reducing metabolic activity and by extension maintaining the dormancy of NWR seeds for longer periods during storage. This could be achieved through the application of exogenous signaling molecules. For seeds of *Panax notoginseng*, which are recalcitrant and have morphophysiological dormancy, exogenous application of abscisic acid (ABA) was able to prolong dormancy and slow embryo development (Wang *et al*. 2023). In addition, nitric oxide has been shown to reduce respiration through the inhibition of the cytochrome pathway (Yamasaki *et al*. 2001; Zottini *et al*. 2002). The reduction of metabolic activity may also be attained through colder storage temperatures than were measured in this study. Previous research has demonstrated that NWR can be stored for at least 26 weeks in non-submerged storage at −3.5°C without killing the seed, though submerged conditions were necessary to break dormancy following this storage type (Kovach and Bradford 1992b; McGilp *et al*. 2020). However, longer term studies are needed to determine how long seeds would stay viable under these conditions. In addition to the above recommendations, researchers should consider measuring the ROS production in submerged storage, with and without aeration, as no studies have focused on their production in submerged rather than hydrated or dry storage.

## Acknowledgements and funding

This work was supported by the State of Minnesota, Agricultural Research, Education, Extension and Technology Transfer (AGREETT) program.

## 5 Author contributions

Protocol development and data collection conducted by Alan Mickelson. Data analysis performed by Lillian McGilp. Writing of original draft and editing done by Lillian McGilp and Jennifer Kimball.

## SUPPLEMENTAL TABLES AND FIGURES

**Table S1.** Percent viable seed mean of Northern Wild Rice (NWR; *Zizania palustris*) averaged across replications and subsamples for each treatment, along with Tukey’s separation for treatment x time.

| Treatment* | Post-Harvest Timepoint | Mean Viable Seed (%)** | Standard Error |
| --- | --- | --- | --- |
| SC | 4 months | 96.67 | 0.96 |
| BC | 4 months | 97.22 | 0.56 |
| WWC | 4 months | 95.00 | 2.55 |
| MWC | 4 months | 97.22 | 1.47 |
| ABC | 4 months | 95.56 | 0.56 |
| AWC | 4 months | 97.22 | 0.56 |
| 2SS | 4 months | 97.78 | 1.11 |
| SC | 9 months | 87.78 | 1.47 |
| BC | 9 months | 93.33 | 0.00 |
| WWC | 9 months | 85.56 | 3.89 |
| MWC | 9 months | 92.78 | 1.47 |
| ABC | 9 months | 64.44 | 0.56 |
| AWC | 9 months | 80.56 | 1.47 |
| 2SS | 9 months | 96.11 | 1.11 |
| SC | 16 months | 88.62 | 3.79 |
| BC | 16 months | 86.71 | 3.78 |
| WWC | 16 months | 51.99 | 14.77 |
| MWC | 16 months | 76.02 | 3.98 |
| ABC | 16 months | 43.05 | 2.73 |
| AWC | 16 months | 36.22 | 7.15 |
| 2SS | 16 months | 75.89 | 3.32 |
| SC | 21 months | 66.17 | 4.75 |
| BC | 21 months | 72.97 | 14.00 |
| WWC | 21 months | 35.59 | 17.29 |
| MWC | 21 months | 62.25 | 7.00 |
| ABC | 21 months | 21.91 | 4.12 |
| AWC | 21 months | 21.29 | 4.43 |
| 2SS | 21 months | 26.44 | 10.40 |
| SC | 28 months | 37.20 | 8.16 |
| BC | 28 months | 28.87 | 4.54 |
| WWC | 28 months | NA | NA |
| MWC | 28 months | 57.63 | 16.20 |
| ABC | 28 months | 28.47 | 10.94 |
| AWC | 28 months | NA | NA |
| 2SS | 28 months | 11.93 | 6.09 |

\*Treatments: SC = standard control; BC = bucket control; WWC= weekly water change; MWC= monthly water change; ABC = aerated, AWC = aerated and weekly water change, 2SS = two stage storage
\*\*Mean Seed Viability (%) was calculated as the %Germinated seed + %Non-Germinated tetrazolium-stained seed

**Supplemental Figure 1.**
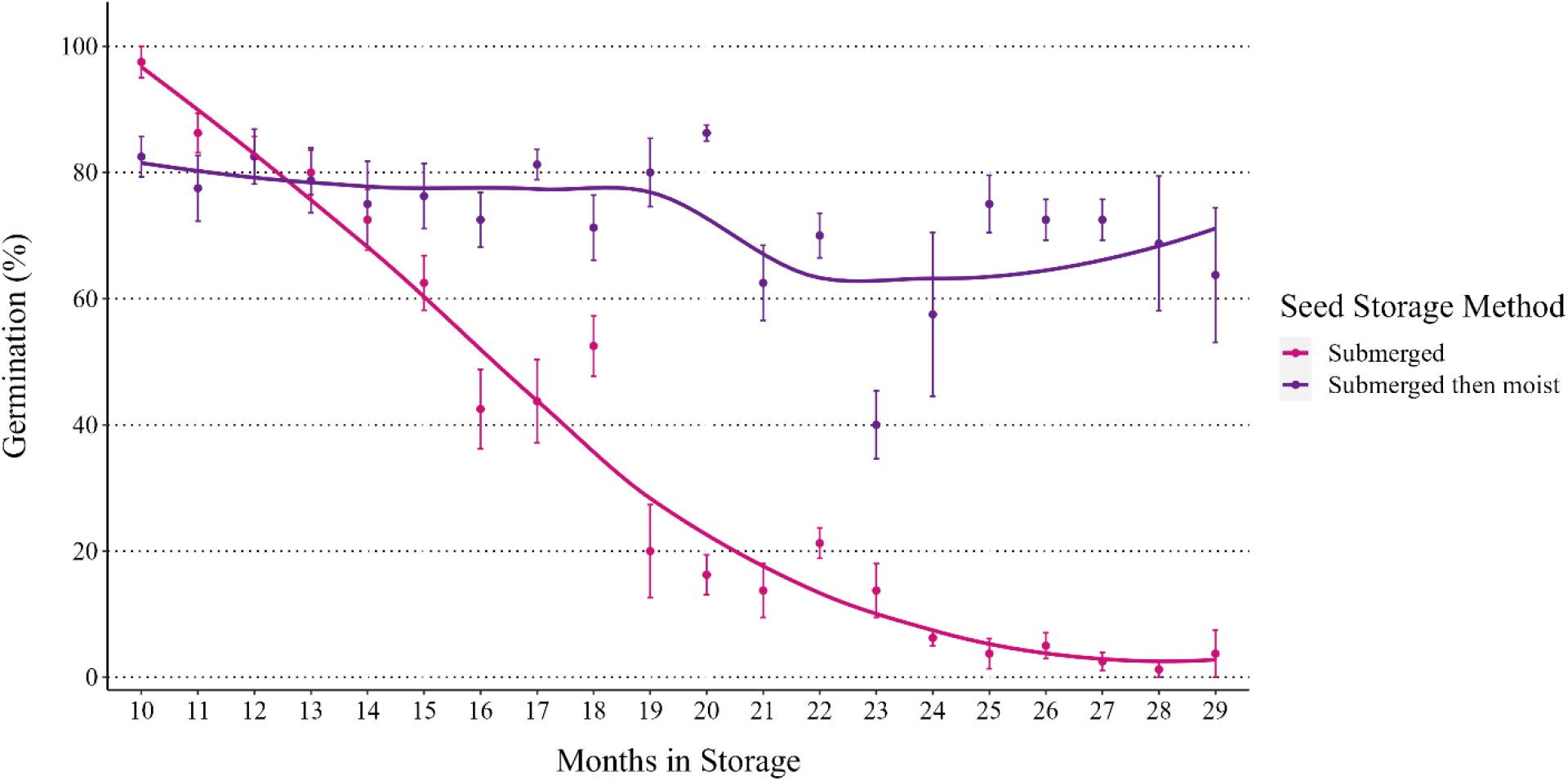
Percent viable seed of Cultivated Northern Wild Rice (NWR; *Zizania palustris*) over the span of 8-29 months in storage. After harvest in September of 2018, seed was stored in large burlap bags in a drainage ditch of a cultivated NWR grower in Waskish, MN until May of 2019 when seed was placed moist but not submerged in standard NWR storage conditions (1-3 □ in the dark).

